# Composition variation of agarwood-associated microbial communities from *Aquilaria sinensis*

**DOI:** 10.1101/583393

**Authors:** Qigui Mo, Chenyang Fan, Gao Zhou, Huiying Fu, Youwei Wang

## Abstract

Agarwood, derived from *Aquilaria sinensis* and *Aquilaria malaccensis*, is of medicinal and ecological value and religious importance as incense. The existing imbalance between short supply and increasing demand of this product remains to be solved. Thus, the biologically artificial agarwood-inducing methods commonly called whole-tree agarwood-inducing techniques (agar-wit) have been established to dramatically improve agarwood yield within a short period. However, several studies reported a lower content of ethanol-soluble extractive in the agar-wit agarwood than in the natural agarwood. To further understand the role of microorganisms in agarwood formation, we investigated and contrasted the endophytic bacteria and fungi between different types of agarwood from *A. sinensis* through high-throughput sequencing. Results showed that the same dominant phyla of bacteria consisting of *Proteobacteria*, *Actinobacteria*, and *Acidobacteria* were shared by the natural agarwood and agar-wit agarwood. Meanwhile, *Ascomycota* and *Basidiomycota* constituted the similar dominant fungal phyla of these two kinds of agarwood. However, the principal microbial communities at the genus or order level evidently varied from natural agarwood to agar-wit agarwood. Moreover, the bacterial communities are closely connected with terpenoid and carbohydrate metabolism, which indicated that the bacterial communities also play a vital role in agarwood formation. In conclusion, the higher concentrated abundance of the dominant microbial communities in agar-wit agarwood than in natural agarwood may promote agarwood formation, however, the low evenness of microbial communities also lowers the content of ethanol-soluble extractive.

**IMPORTANCE:** Agarwood has become an indispensable product in modern life because of medicinal value, ecological and religious importance as incense. Nevertheless, the enormous demand for agarwood markedly exceeds the supply because of the dramatically declining population of genus *Aquilaria*. Agarwood formation occurring slowly and infrequently in a natural environment, so various artificial techniques were developed to promote the formation of agarwood, such as the physical methods and chemical methods. However, these techniques still are insufficient to compensate for the agarwood shortage. In this case, a novel biological method called the whole-tree agarwood-inducing technique (Agar-wit) induces agarwood production. However, several studies have shown that agarwood harvested from biological technology contains lower content of ethanol-soluble extractive compared with natural agarwood. So to further expose and understand the endophytic bacteria and fungi in agarwood formation is important for the improvement of biological method.

## INTRODUCTION

*Aquilaria sinensis* (Lour.) Gilg (*Thymelaeaceae*) is mainly distributed in the provinces of Hainan, Guangdong, Fujian, and Yunnan in China (1, 2), and *Aquilaria malaccensis* Lam is native to Malaysia (3). These plants are sources of agarwood, which is the dark resinous wood harvested from the tree (China Pharmacopoeia 2015, first section). Agarwood is highly prized for its aphrodisiac, sedative, cardiotonic, and carminative effects and its ability to relieve gastric problems, coughs, rheumatism, and fever (4). Agarwood (Chen Xiang in China), used as a traditional Chinese medicine to dispel damp toxin, was first documented in Mingyi Bielu in the Wei and Jin Dynasties. Traditional use was complementally recorded in Bencao Gangmu in the Ming Dynasties. Recently, many chemical ingredients of agarwood have revealed significant anticancer (5) and anti-inflammatory activities (6). Moreover, agarwood has been used as incense for centuries in Buddhist, Hindi, and Islamic ceremonies (4). DingWei, a famous writer in the Song Dynasty, provided crucial contributions to the development of agarwood as incense. In Tianxiang Zhuan, he was the first to classify agarwood into four grades and 12 shape categories depending on the fragrance, appearance, and formation. In a word, agarwood has become an indispensable product in modern life.

Nevertheless, the enormous demand for agarwood markedly exceeds its supply because of the dramatically declining population of the genus *Aquilaria* caused by illegal overexploitation (7). This genus has been listed in Appendix II of the Convention on International Trade in Endangered Species of Wild Fauna and Flora since 2004 (1). The factors mentioned above explain the price inflation of agarwood products to US$ 100,000/kg for superior, pure material (4). Natural agarwood forms around trunk wounds, particularly external wounds caused by lightning strikes, animal attack, which easily induce the wounds to infect with microorganisms (8). As agarwood formation occurs slowly and infrequently in the natural environment, various artificial techniques have been developed to promote agarwood formation. Such techniques include physical methods where tree trunks and branches are wounded using axes, fire, knives, or nails (9) and chemical methods where the wounds are intentionally treated with some chemical reagents initially (e.g., inorganic salt, acids, and phytohormone) (10).

However, these techniques are insufficient to compensate for the agarwood shortage. In this case, a novel biological method called the whole-tree agarwood-inducing technique (agar-wit) induces agarwood production by inoculating certain elicitors into the xylem part of *Aquilaria* trees through simple transfusion sets (12). The prepared inducer is then transported throughout the whole tree gradually by transpiration force and initiates a series of defensive responses to produce resin (12). A remarkable biological inoculation is the fungal inoculum, which is mainly fermentation liquid of fungi, such as *Menanotus flavolives* (13), *Botryosphaeria dothidea* (14), and *Lasiodiplodia theobromae* (12). As reported, such inoculum is capable of stimulating agarwood formation. Given the fungal mobility and plant transpiration, agarwood may form on the trunk and branches of the entire tree, causing a far higher production of agarwood at a lower cost of labor and time than that induced by previous artificial techniques. However, several studies have shown that agarwood harvested from biological technology contains lower content of ethanol-soluble extractive compared with natural agarwood (15–17). Zou (18) considered the possible role of endophytic bacteria in agarwood formation, but failed to determine the bacterial community structure by PCR–DGGE methods. Thus, the bacterial and fungal communities of agarwood from *A. sinensis* must be investigated to refine the current agar-wit technique.

In this study, the major bacterial and fungal communities of natural agarwood and agar-wit agarwood were investigated using high-throughput sequencing without the prepared isolation and cultures. The variation of dominant microbial communities between these two agarwood types was analyzed. Functional profiles were also used to investigate the role of bacterial communities in the metabolic activity of agarwood formation. Finally, the relationship between the variation of dominant microbial communities and agarwood types was characterized.

## RESULTS

General analyses. After data filtration, a total of 64474 and 37850 high-quality sequences of all samples were obtained for the fungal primer SSU0817F/1196R and bacterial primers 799F/1392R and 799F/1193R, respectively. Then, these effective sequences were clustered to OTUs with 97% similarity, which is a common parameter for describing microbial community structure (19, 20). After the chimeric, mitochondrial, and chloroplast sequences were removed, the OTU table generated 30 OTUs and 185 OTUs for fungal and bacterial communities, respectively. These OTU tables were used for downstream analyses.

First, rarefaction curves were applied to evaluate and compare data from the current study (Fig. S1). With increasing read number, the curves tended to stabilize and be saturated for both bacterial and fungal taxonomies; this finding suggested that the sequenced numbers were adequate to reflect the vast majority of microbial types, and the structures of bacterial and fungal communities associated with different agarwood samples could be reasonably characterized by a sufficient saturated OTU number (21).

### Bacterial and fungal diversity analyses

Chao and Shannon indices were related to microbial communities’ richness and diversity and are usually used to preliminarily compare the different types of samples (22–24). In our study, the Chao and Shannon indices of bacteria (Fig. 1a and 1b) in natural agarwood samples were higher than those in agar-wit samples, and the same trend was found in fungal communities (Fig. 1c and 1d). These results indicated that the microbial types in natural agarwood, including bacteria and fungi, were more abundant and complicated than those in the agar-wit samples. In both natural agarwood and agar-wit samples, the values of Chao and Shannon indices of bacterial types (Fig. 1a and 1b) were significantly higher than those of fungal types (Fig. 1c and 1d), which suggested that the bacterial communities showed higher abundance than the fungal communities in agarwood samples from two different production methods.

**FIG 1.**
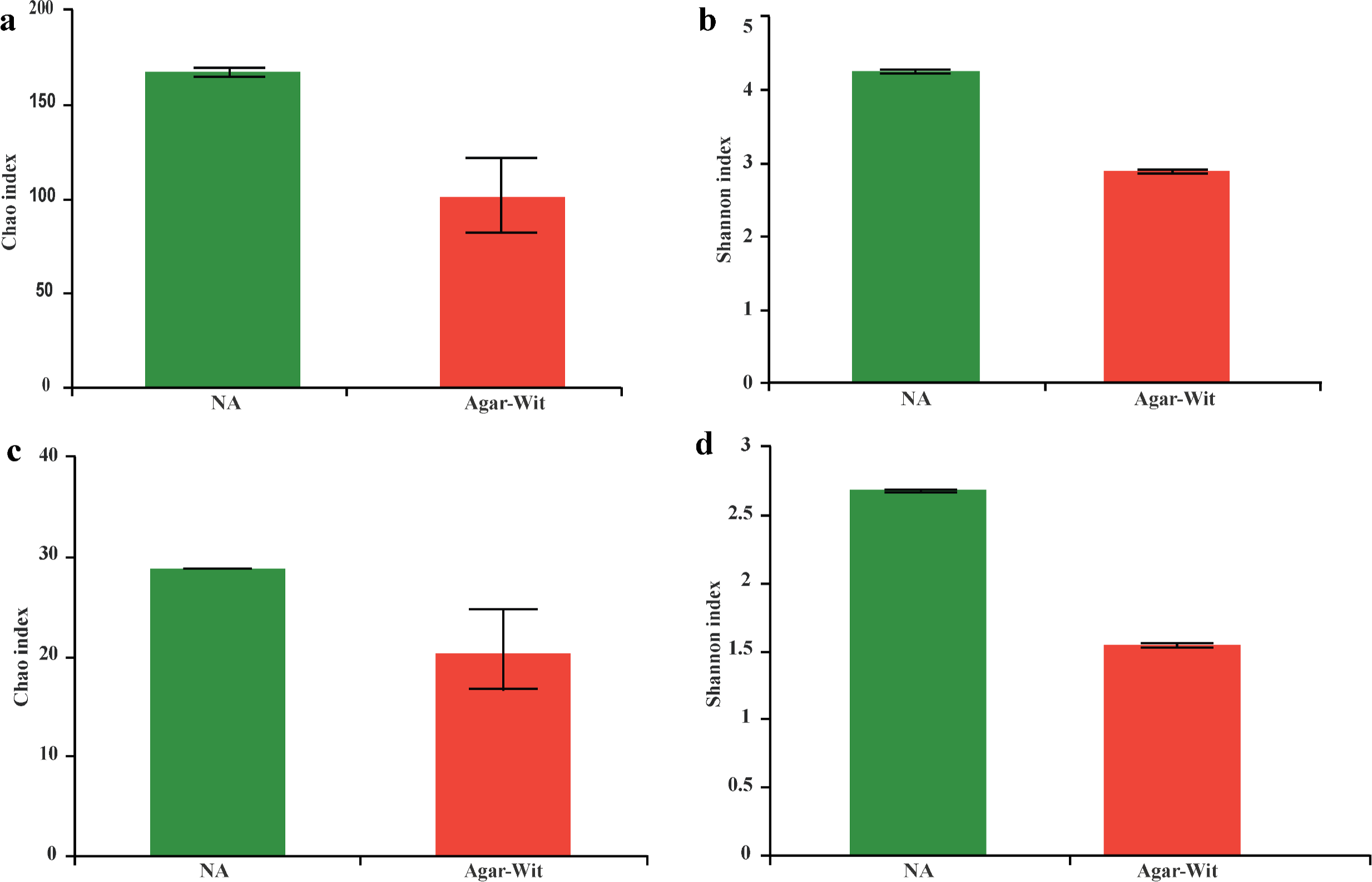
Community richness and community diversity indices at a 97% identity threshold of the natural agarwood (NA) and agar-wit agarwood (Agar-Wit). a and c are the Chao indices of bacterial and fungal communities, respectively. b and d are the Shannon indices of bacterial and fungal communities, respectively. The values are presented as means ± SD.

### Cross-OTU comparisons in natural agarwood and agar-wit samples

To determine the relation between these two kinds of samples, we conducted cross-OTU comparison by using Venn diagrams (25). The overlaps were present in two different types of microorganisms between natural and agar-wit agarwood (Fig. 2a and 2b). The bacterial and fungal OTUs enriched in the natural agarwood successfully colonized the agar-wit agarwood, as 77 out of 185 OTUs of bacteria included in natural agarwood were also enriched in the agar-wit agarwood (Fig. 2a), and the fungal OTUs in the agar-wit agarwood were almost completely replicated from natural agarwood (Fig. 2b). These results indicated that the bacterial species in agarwood obtained from whole-tree agarwood-induction technology were stabilized and only slightly varied relative to that in natural agarwood.

**FIG 2.**
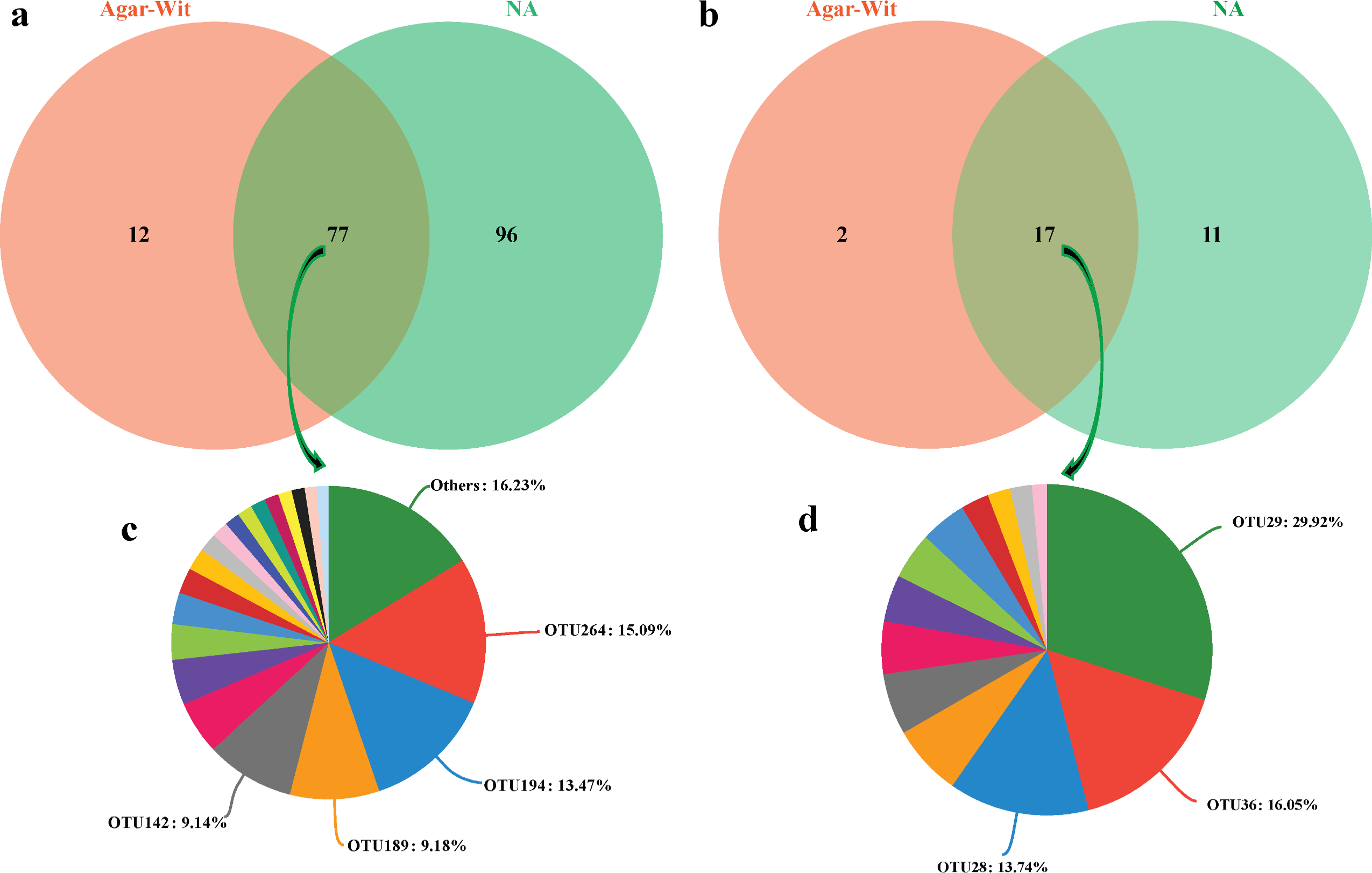
Venn diagrams depicting the shared or unique OTUs for natural agarwood (NA) and agar-wit agarwood (Agar-Wit). The OTU abundance with < 1% are combined and defined as others. a and b are bacterial and fungal OTUs, respectively. c and d are the main shared OTUs in bacterial and fungal types, respectively.

A total of 77 and 17 OTUs were clustered to the common region, which were all shared by two different agarwood samples for bacterial and fungal types, respectively (Fig. 2c and 2d). A total of 77 OTUs for bacteria mainly consisted of *Proteobacteria* and *Actinobacteria* (Fig. 3a), and 17 OTUs for fungi were mainly composed of *Ascomycota* (Fig. 3b). These same OTUs were enriched in two different agarwood samples with different abundances. The majority of *Proteobacteria* bacterial OTUs were allocated to *Burkholderiaceae* (15.09%) and *Sphingomonadaceae* (13.47%), whereas most of the *Actinobacteria* bacterial OTUs corresponded to *Microbacteriaceae* (9.18%) and *Solirubrobacterales* (9.14%). The *Ascomycota* fungal OTUs contained *Coniochaetaceae* (29.92%) and *Sordariomycetes* (29.79%, OTUs 36 and 28). These results indicated that the shared microbial communities by natural agarwood and agar-wit agarwood were concentrated on several types.

**FIG 3.**
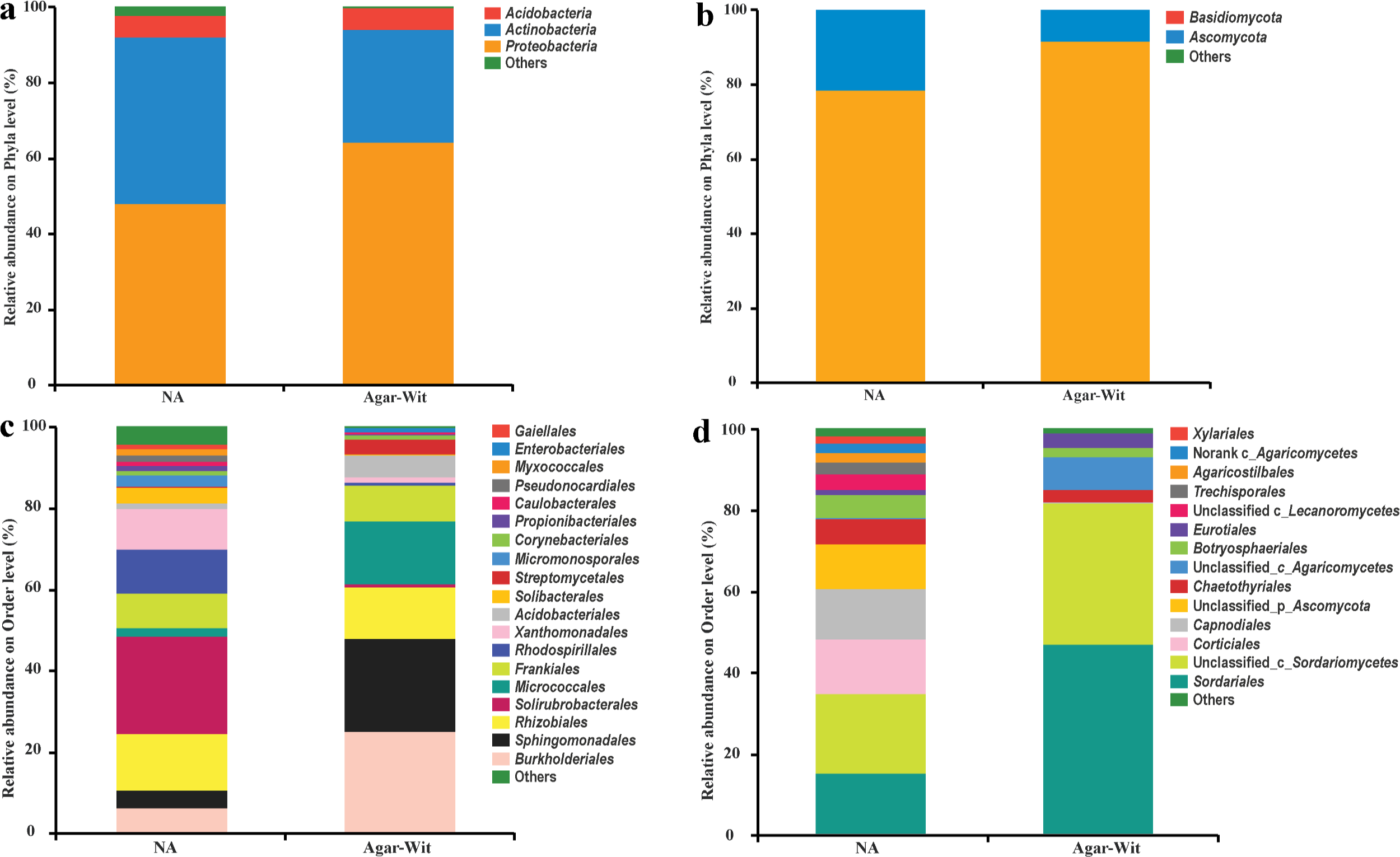
Bacterial and fungal compositions of natural agarwood (NA) and agar-wit agarwood (Agar-Wit) on the phyla (a, b) and order (c, d) levels. a and c are the bacterial communities, and b and d are the fungal communities. Others include the few sequences with < 1% relative abundance.

### Microbial community composition in natural agarwood and agar-wit agarwood

In this study, we investigated the bacteria at different taxonomic levels, and taxonomies with < 1% abundance were defined as others (Fig. 3a and 3c). The dominant phyla consisted of *Proteobacteria*, *Actinobacteria*, and *Acidobacteria*, which constituted the main sections of the bacterial community structure of natural agarwood and agar-wit agarwood (Fig. 3a). The agar-wit technology distinctly caused an increasing ratio of *Proteobacteria* in the agar-wit agarwood from 47.87% to 64.04% compared with natural agarwood. Inversely, the relative abundance of *Actinobacteria* decreased from 44.08% of natural agarwood to 30.06% of agar-wit agarwood and showed a reverse trend (Fig. 3a). Similar to *Arabidopsis* (26) and rice (25), an increased pool of *Proteobacteria* was found in the bacterial communities. However, the difference was that the bacterial community composition was simpler in agarwood, and the three bacterial phyla above achieved a total relative abundance of more than 97.60%. The simpler bacterial community composition of agarwood might be due to the lack of soil microorganisms that could influence each other directly compared with *Arabidopsis* and rice. More detailed variation in relative phylum abundance between two different samples could be embodied by a more specific order level (Fig. 3c). Compared with natural agarwood, the substantially increasing ratio of *Burkholderiales* and *Sphingomonadales* and the sharply decreasing ratio of *Solirubrobacterales* in agar-wit agarwood samples were the main reasons for the change in the phyla of *Proteobacteria* and *Actinobacteria* (Fig. 3c). Interestingly, *Xanthomonadales*, which belongs to the phylum *Proteobacteria*, sharply decreased in agar-wit agarwood than in natural agarwood, which was not observed with the phylum *Proteobacteria*. Similarly, the variation tendency of *Micrococcales* was vastly increasing and thus opposite to that of phylum *Actinobacteria* (Fig. 3c). The variation in relative abundance of *Xanthomonadales* and *Micrococcales* may be due to the cooperation and competition among the different microbial types, which have been thoroughly investigated and utilized (27, 28). More details were showed in Table S1. 65.95% OTUs were assigned to genus or species.

At present, fungi are considered as the most important microbial factor promoting agarwood formation. Therefore, studies on fungal communities existing in the resinous region of *A. sinensis* emerged in succession over the years (14, 18, 29, 30). In our study, high-throughput sequencing without the prepared isolation and cultures was applied to characterize the fungal community composition in natural agarwood and agar-wit agarwood. The chloroplast, *Amoebozoa* OTUs, and other not belonged to fungi were filtrated, and then 30 OTUs, which included 1 singleton, were obtained. Based on the results, *Ascomycota* and *Basidiomycota* constituted the main fungal community composition with a total relative abundance of more than 99.89% (Fig. 3a and 3b). Consistent with Premalatha’s (30) report, the phylum *Ascomycota* occupied the majority of sequence numbers with 78.55% relative abundance in natural agarwood and 91.68% relative abundance in agar-wit agarwood (Fig. 3b). On the contrary, the relative abundance of *Basidiomycota* was 21.34% in natural agarwood, which sharply decreased to 8.32% in agar-wit agarwood (Fig. 3b). At the taxonomic order level, the proportion of phylum *Ascomycota* in agar-wit agarwood indicated a rise relative to natural agarwood owing to the increasing percentage of *Sordariales* and another unclassified order. Actually, these two fungal types on the order level belonged to the same class, *Sordariomycetes* (Fig. 3d). Similar to other bacterial communities, the antagonism could manage the relative abundance of some fungal types. The sharply decreasing *Capnodiales* ratio in agar-wit agarwood was mainly attributed to the lost vivosphere, which was compressed by other dominant fungal types. The final taxons of all fungal OTUs were showed in Table 1. Only 8 OTUs were assignable to genus or species. To supplement this, 10 fungal strains were isolated and identified, and all strains were assigned to genus or species. 6 strains belonged to *Sordariomycetes* (Table S2), which also were the most abundant of microbial types in the culture-independent approach (Fig. S2). *L. theobromae*, a strain usually used in whole-tree agarwood-inducing technique, was isolated from both the natural agarwood and agar-wit agarwood.

**TABLE 1.**
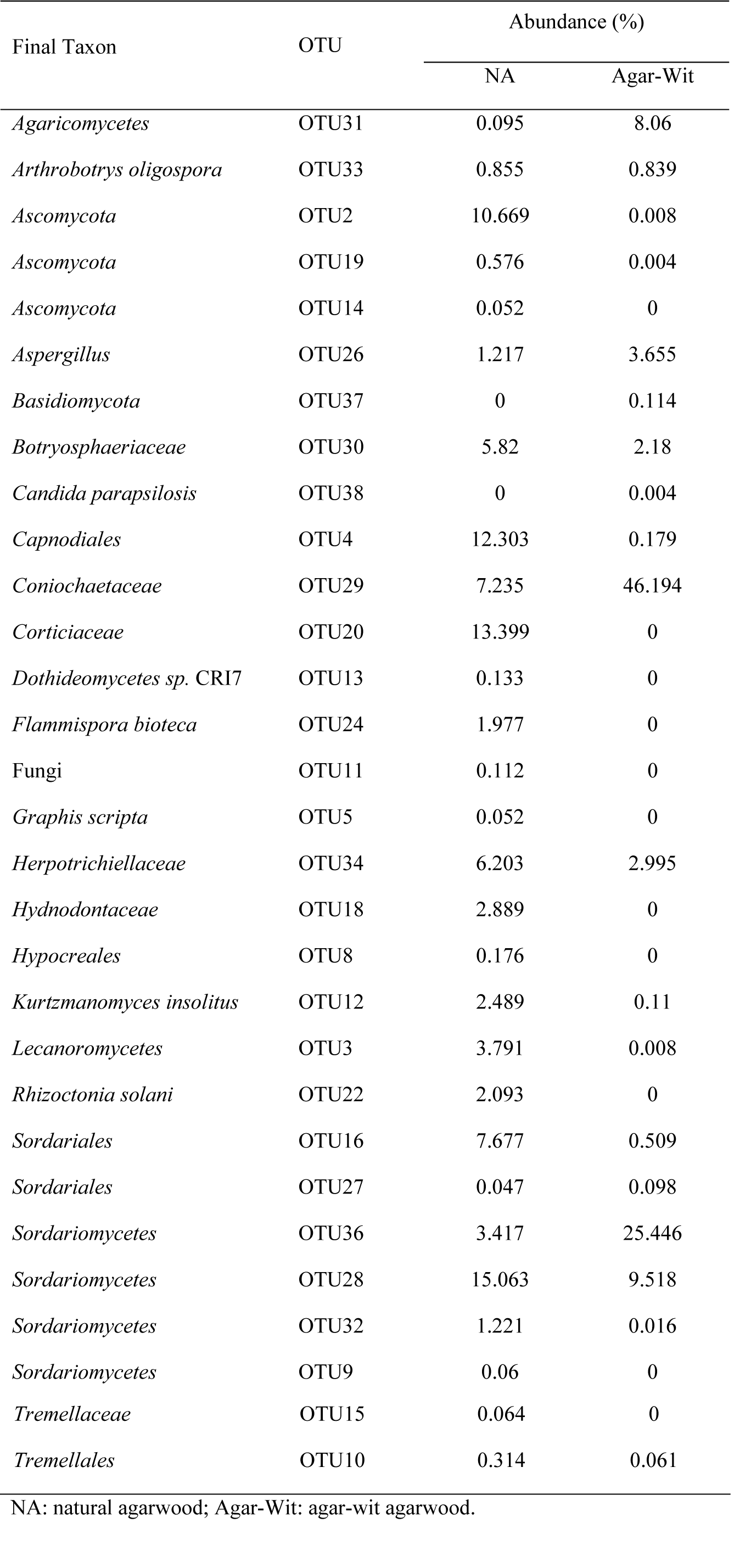
The final taxon of all fungal OTUs detected from high-throughput sequencing.

### Comparative analyses between natural agarwood and agar-wit agarwood

In the heatmap of bacterial and fungal distributions, we used color intensity to compare the top 50 most abundant genera present in the natural agarwood and agar-wit agarwood (Fig. 4). Lower evenness was found in the bacterial communities of the agar-wit agarwood than in the natural agarwood (Fig. 4a). The bacterial types in the agar-wit agarwood focused on several genus levels, such as *Gryllotalpicola*, *Burkholderia*-*Paraburkholderia*, *Sphingomonas*, and *Acidothermus* (Fig. 4a). A same trend was found in the fungal communities of the natural and agar-wit agarwood (Fig. 4b). In particular, the distribution revealed a highly concentrated relative abundance of *Sordarionmycetes* (Fig. S3).

**FIG 4.**
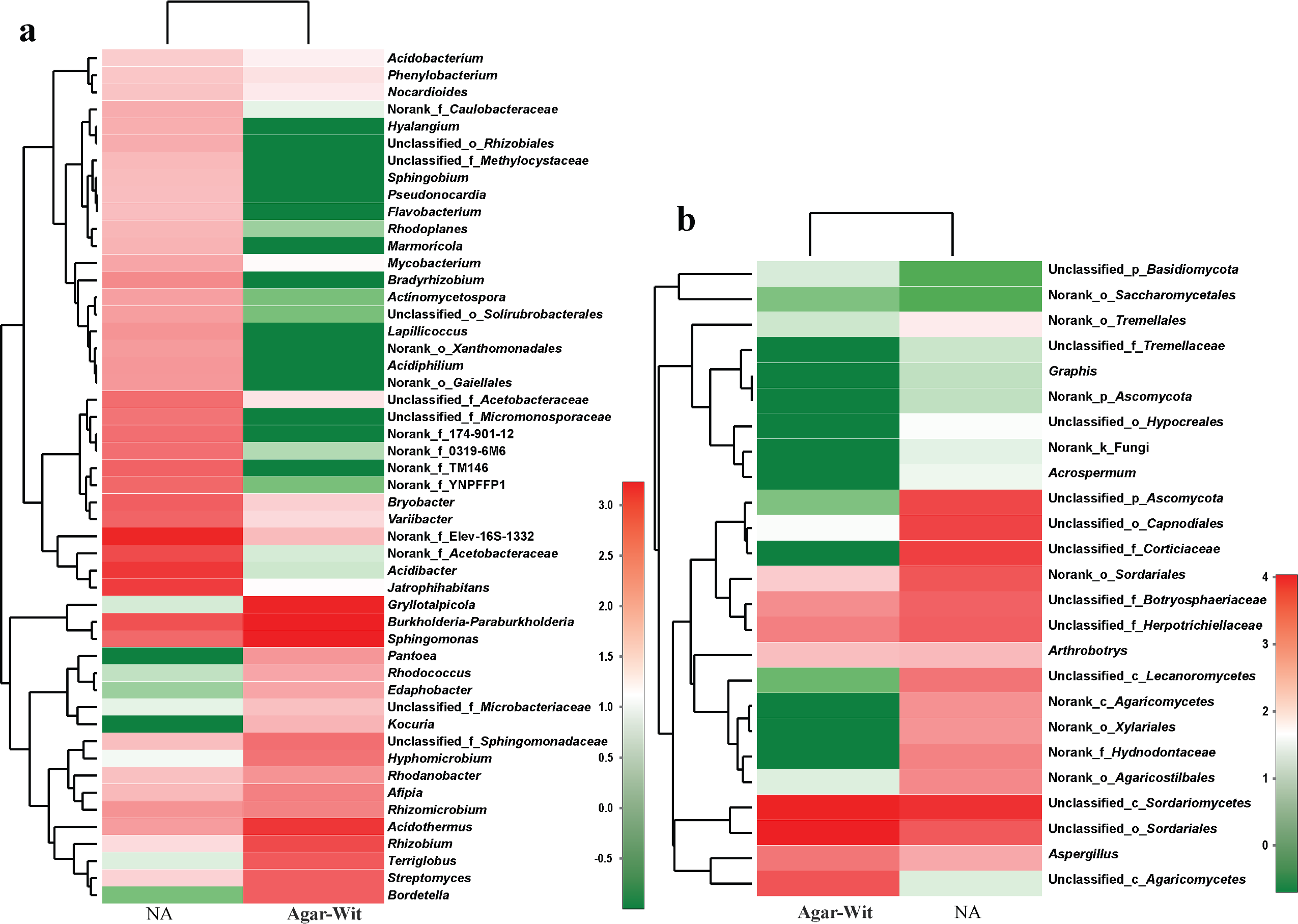
Heatmap of the bacterial (a) and fungal (b) distributions of the top 50 most abundant genera presented in the natural agarwood (NA) and agar-wit agarwood (Agar-Wit). The relative abundances of bacterial and fungal genera are indicated by color intensity. In the current study, the total fungal community types of these two samples were 25.

The highly concentrated relative abundance of some microbial communities caused variation in the dominant microbial communities between natural agarwood and agar-wit agarwood (Fig. 5). The main dominant bacterial communities at the genus level contained *Jatrophihabitans*, *Acidibacter*, and an unclassified genus belonging to the family *Elev-16S-1332* in natural agarwood (Fig. 5a and Fig. S4). However, the highly abundant population was changed to *Burkholderia-Paraburkholderia*, *Sphingomonas*, *Gryllotalpicola*, and *Acidothermus* in agar-wit agarwood. Actually, the relative abundance of the bacterial communities at the genus level tended to be thoroughly distributed in natural agarwood, which showed a higher evenness in bacterial communities than that in agar-wit agarwood. A variation in the dominant microbial communities was also found within fungal types on the order level between natural agarwood and agar-wit agarwood (Fig. 5b). The dominant fungal communities changed from *Sordariomycetes*, *Agaricomycetes*, and *Dothideomycetes* in natural agarwood to *Sordariomycetes* in agar-wit agarwood.

**FIG 5.**
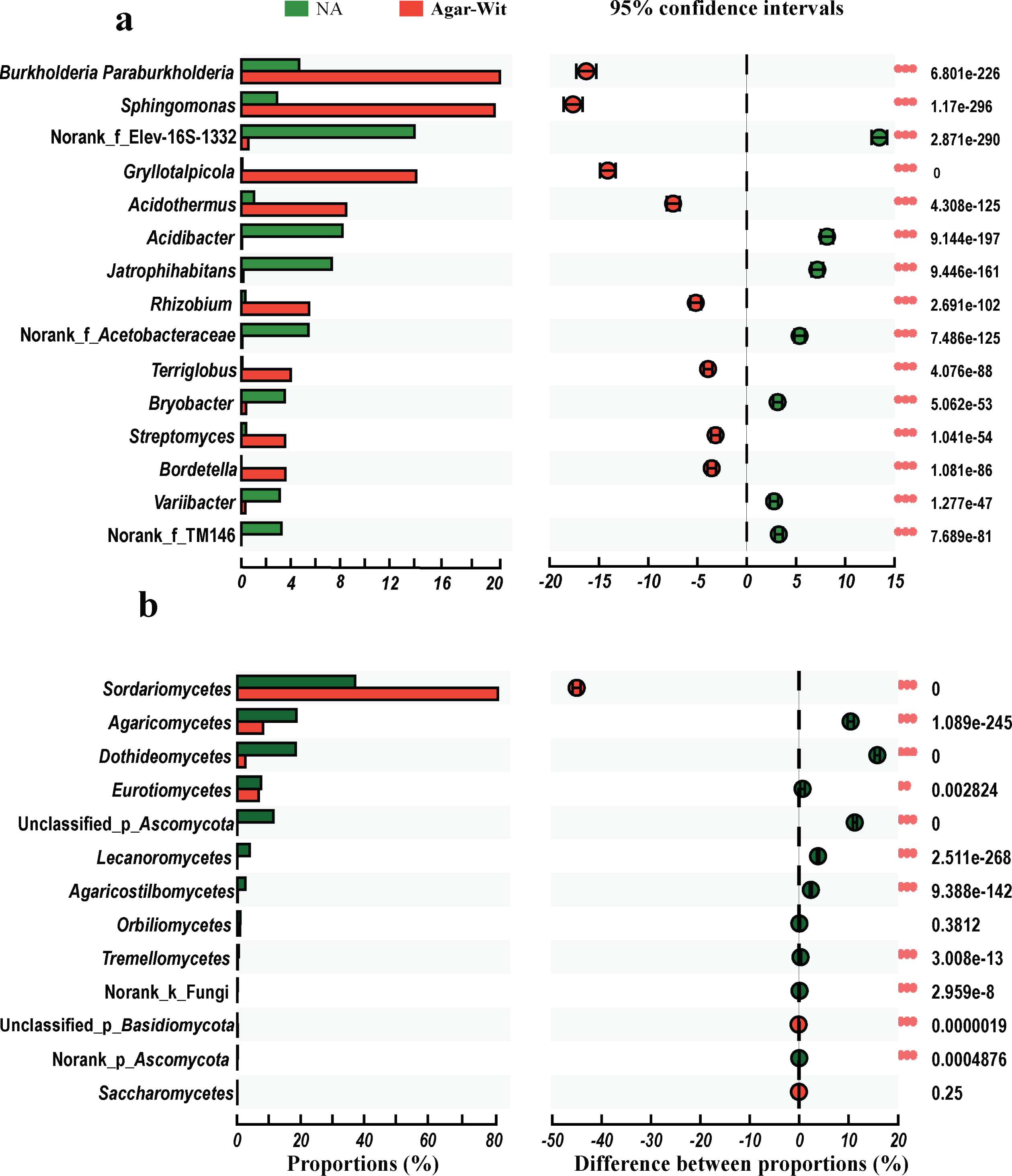
Taxon-level analysis of individual genera or orders altered between the natural agarwood (NA) and agar-wit agarwood (Agar-Wit) by Fisher’s exact test. a: Bacterial communities at the genus level; b: fungal communities at the order level. The right graph indicates the altered percentage of species abundance in the confidence interval. **0.001 < *p* < 0.01, \*\*\**p* < 0.001.

### Bacterial functional profile

The PICRUSt tool has been used to predict the functional profile of the 16S rRNA gene and achieves a high correspondence with the reference genome across several microbial types (31, 32). This method thoroughly compensates for the weakness of 16S rRNA, which cannot directly provide a functional profile (31). By annotating the Cluster of Orthologous Group (COG) function, we obtained 25 functional classifications, in which general function prediction only and function unknown were the two largest clusters. This result was very similar to a previous study (33). After removing the two above-mentioned functional classifications that tentatively held no connection with agarwood formation, we compared the top 18 most abundant functional classifications (Fig. 6a). As a result, two annotated functions related to agarwood formation, “Carbohydrate transport and metabolism” and “Secondary metabolites biosynthesis, transport and catabolism,” shared a high relative abundance in the agarwood samples. These two metabolic activities were more active in agar-wit agarwood than in natural agarwood (Fig. 6a). After annotating the Kyoto Encyclopedia of Genes and Genomes (KEGG) function, we obtained above 52.64% relative abundance of metabolic pathways (Fig. 6b). Terpenoid and polyketide metabolism is an important biosynthetic pathway of sesquiterpenes in this study, and besides, carbohydrate metabolism also showed a prominent abundance in two types of agarwood (Fig. 6c). Most of the COG and KEGG functions were higher in agar-wit agarwood than in natural agarwood probably because of the variation in the dominant bacterial communities.

**FIG 6.**
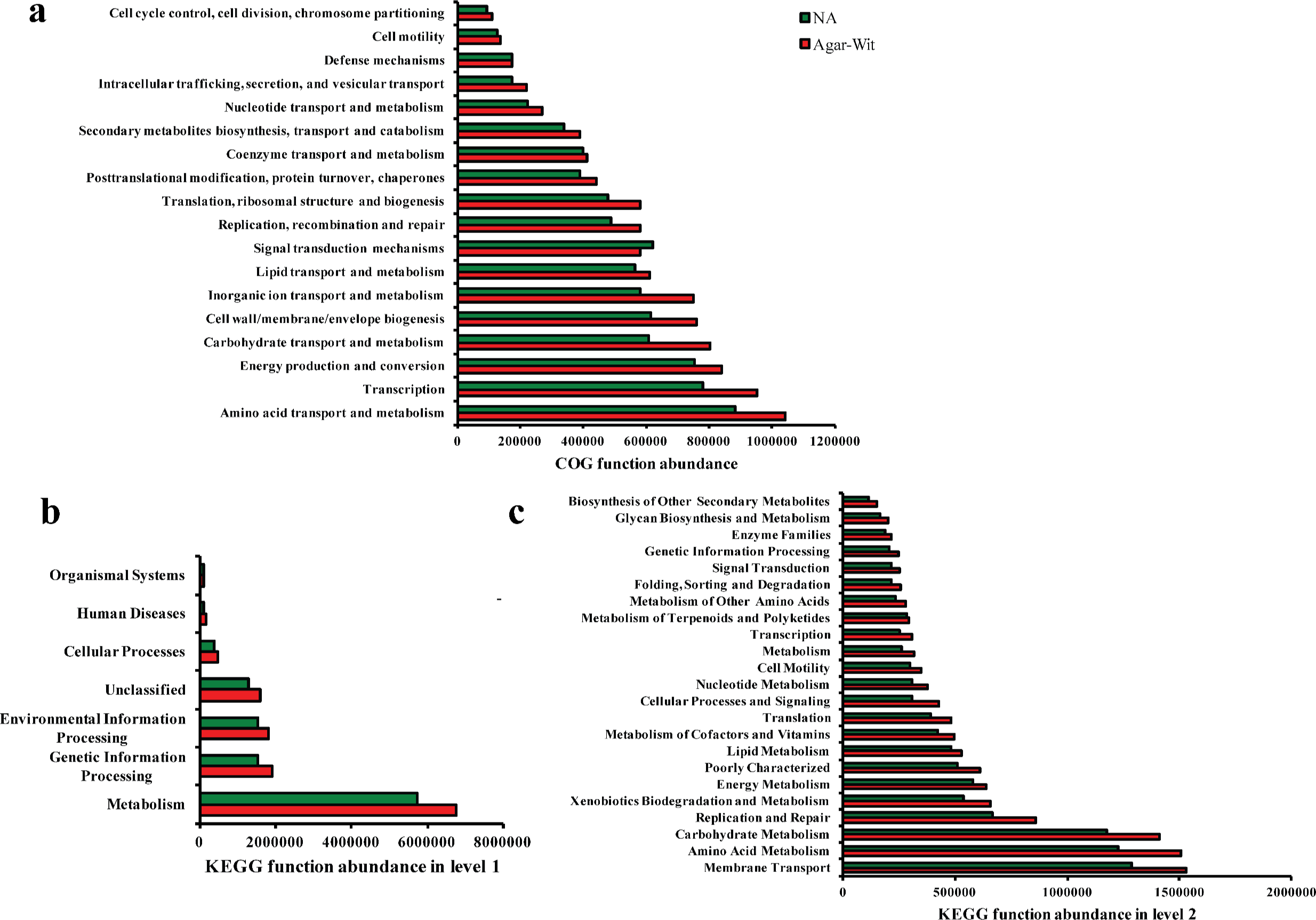
Abundance and functional predictions of bacterial communities of the natural agarwood (NA) and agar-wit agarwood (Agar-Wit). a: COG functional predictions of OTUs. b and c: KEGG functional predictions of OTUs by using the PICRUSt tool in category levels 1 and level 2, respectively.

## DISSCUSSION

Agarwood are more and more popular in our daily life as medicine and incense nowadays. Majority of techniques, mainly including physical, chemical, and biological methods have been established to promote agarwood yield (9–11). Expressly, the biological technique, which mainly inoculated fungal fermentation liquid into *A. sinensis* trees, has been considered as the most promising method to obtain a far higher production of agarwood at a lower cost of labor and time (12–14). So previous studies on microbiomes from *A. sinensis* mainly focused on fungal communities across the white wood and resinous region (14, 29, 30). In the present study, we successfully characterized the bacterial and fungal community structure in the natural and agar-wit agarwood through high-throughput sequencing. We also found that bacterial communities were deeply involved in agarwood formation.

Actually, bacterial communities are drew less attention compared to fungal communities in agarwood formation. Premalatha and Kalra (30), who proposed that pathogenic fungi are the primary causative factor in agarwood formation, attempted to investigate the impact of endophytic fungi on the resinous and white wood of *A. malaccensis*. Zou (18) considered that endophytic fungi and bacteria were involved in agarwood formation, but failed to reveal the bacterial community structure by PCR– DGGE methods. In this study, we found the higher bacterial communities’ richness and diversity in both types of agarwood than fungal communities (Fig. 1). Remarkably, we detected a high bacterial pool from *A*. *sinensis* by high-throughput sequencing, which was typically ignored in the past.

The diversity of microbial communities between the natural and agar-wit agarwood were different in current study. The microbial communities richness and diversity analyses all showed the higher values in the natural agarwood than agar-wit agarwood (Fig. 1). Venn diagrams indicated that the microbial species existing in agarwood obtained from whole-tree agarwood-induction technology were relatively fixed and only slightly changed relative to that in the natural agarwood (Fig. 2). Although Zou (18) considered that fungal species were significantly abundant in agarwood samples compared with white wood, other sufficient reports shared different ideas. Mohamed (29) found no difference in fungal species between agarwood and white wood. Similarly, Tian et al (14) reported that most of the fungi on the genus level colonized the white wood from resinous sections. Combined with our results, we boldly speculate that the methods to promote agarwood formation, physical, chemical, and microbial infection methods, enhanced the abundance of key microbial species rather than transforming the microbial community structure.

Fungal infection technology promoting agarwood formation is becoming increasingly developed, but this method still has a noticeable deficiency. Several studies proved that the content of ethanol-soluble extractive, a crucial indicator in evaluating agarwood product quality in the Chinese Pharmacopeia (2015), was lower in the agarwood product from agar-wit technology than in the natural agarwood samples (15–17). Therefore, the role of bacterial infection in agarwood formation is increasingly being valued. We successfully characterize the bacterial community composition of agarwood products through high-throughput sequencing, an advanced method that allows the investigation of microbial types without the prepared isolation and cultures (25, 34). Although the richness and diversity of microbial communities and cross-OUT comparison analyses showed the difference of microbial communities in the natural and aga-wit agarwood, these two types of samples shared the same dominant phyla for both bacterial and fungal communities (Fig. 3a and 3b). On order level, the abundances of most microbial categories were different, which might cause the different values of microbial communities’ richness and diversity.

In particular, unlike in most previous studies, the comparative analysis of *A. sinensis* between the white wood and the resinous region was performed to determine the most important microbial communities in agarwood formation (14, 18, 29, 30). In our study, we compared the bacterial and fungal types between the natural agarwood and agar-wit agarwood and found a variation in the dominant microbial communities between these two kinds of agarwood samples. In the cross-OTU comparative analysis, we summarized that the bacterial and fungal species enriched in the natural agarwood successfully colonized the agar-wit agarwood. This result indicated no difference in microbial types between these two kinds of agarwood. In the current study, the primer SSU0817F/1196R, which showed a fungal-specific characteristic, was employed to accurately characterize the fungal communities without any other false-positive results (35). However, the taxonomic identification from this primer was restricted (35). In our study, most of the fungal communities were characterized on the order level or above (Fig. S3). Actually, next-generation sequencing technology (NGS) have many constrains (36). The fungal DNA may not be recovered from all genotypes in the culture-independent approaches (36). Besides, PCR amplification steps may prefer to the bulk DNA extracts, and hinder the identification of some other genotypes (37). Finally, fungal sequences obtained from the culture-independent approaches are short and variable, which restrict the assignment of fungal sequences to genus or species (38). So the culture-independent methods are usually used to reflect the microbial communities, and the culture-dependent methods are used to identify the microbial taxons. These might be the reasons why no genus or species overlapped between the culture-dependent and culture-independent approaches in this study. Despite all these shortcomings, a lower evenness of fungal communities was found in the agar-wit agarwood than in natural agarwood at the genus level (Fig. 4b). Bacterial communities also showed the lower evenness in agar-wit agarwood and focused on several genus levels (Fig. 4a). The lower evenness and higher concentrated relative abundance of microbial communities resulted in variation of the dominant microbial communities between natural agarwood and agar-wit agarwood (Fig. 5). Given the above results, the highly concentrated relative abundance of the dominant microbial communities in agar-wit agarwood might be a key factor in promoting agarwood formation relative to that in natural agarwood. However, the decreased evenness of microbial communities also caused the lower content of ethanol-soluble extractives.

The role of bacterial communities in agarwood formation is unknown, we proved that bacterial communities were deeply involved in agarwood formation in current study. The annotated COG and KEGG functions showed that bacterial communities shared the abundant functions of terpenoid and carbohydrate metabolism in natural and aga-wit agarwood. Sesquiterpenes comprise the main component of agarwood (39). Sesquiterpene synthesis begins with some complex pathways from glycolysis (40, 41). Thus, carbohydrate metabolism with high relative abundance can provide a material for sesquiterpene synthesis in agarwood formation (Fig. 6c). The increased level of terpenoid and carbohydrate metabolism benefits agarwood formation in agar-wit agarwood samples. Several bacteria have been reported to participate in the synthesis of terpenes (42–45). In this study, an *Mycobacterium* OTU and three *Streptomyces* OTUs were identified (Table S1), which might be the probable contributors for the COG and KEGG functions of terpenoid and carbohydrate metabolism. In this section, the results of the functional profile of the 16S rDNA gene demonstrated that bacterial communities also played a critical role in agarwood formation.

In summary, we successfully characterized the bacterial and fungal community composition in natural agarwood and agar-wit agarwood. In these two kinds of agarwood products, the dominant phyla of bacteria consisted of *Proteobacteria*, *Actinobacteria*, and *Acidobacteria*. Meanwhile, the same predominant fungal phyla included *Ascomycota* and *Basidiomycota*. The variation of the dominant microbial communities between natural agarwood and agar-wit agarwood has been described at the taxonomic level of genera or order. The functional profile predicted using the PICRUSt tool indicated that bacterial communities were deeply involved in agarwood formation. We inferred that the highly concentrated relative abundance of the dominant microbial communities in agar-wit agarwood can more effectively promote agarwood formation than that in natural agarwood. However, the low evenness of microbial communities also results in the low content of ethanol-soluble extractive. We believe that this study will make it easier to understand the role of bacterial communities in agarwood formation.

## MATERIALS AND METHODS

### Agarwood samples

Two kinds of agarwood samples were collected from mature *A. sinensis* (Lour.) Gilg trees in Haikou, China (E109°27′26.08″, N19°42′20.02″), in June 24, 2017. Natural agarwood samples were cut off at 2–3 cm thickness from the exposed section where agarwood formed naturally. Agar-wit samples were then collected from the bottom of the trunk into which fungal inducers were injected 24 months earlier. All samples were cut with a sterilized knife or saw, placed in clean sealable bags immediately, and transported back to the laboratory under 4 °C. Excess white wood was removed, and the remaining agarwood was surface sterilized with 75% ethanol and finally stored in an ultra-low-temperature freezer (Haier biomedical, Qingdao, China) for endophytic microbial diversity analysis.

### DNA extraction

A series of experiments, including DNA extraction, PCR amplification, and pyrosequencing, were carried out on the agarwood samples by Majorbio Bio-Pharm Technology Co., Ltd. (Shanghai, China). Briefly, agarwood samples were separately milled into powder under liquid nitrogen. Total DNA was extracted from 0.5 g powdered agarwood in accordance with a previously described CTAB (Cetyltrimethylammonium Ammonium Bromide) method (30). Total DNA was quantified by agarose gel electrophoresis and then used for PCR amplification.

### PCR amplification of the V5–V7 region of the fungal 18S and bacterial 16S rRNA genes

The V5–V7 region of the 18S rRNA fungal-specific primers (SSU0817F: 5′-TTAGCATGGAATAATRRAATAGGA-3′ and 1196R: 5′-TCTGGACCTGGTGAGTTTCC-3′) were designed for endophytic fungal diversity analysis (35). Meanwhile, to accurately assess the endophytic bacterial community, two sets of amplification primers were designed. The first one was 799F/1392R (799F: 5′-AACMGGATTAGATACCCKG-3′ and 1392R: 5′-ACGGGCGGTGTGTRC-3′), and the second was 799F/1193R (799F: 5′-AACMGGATTAGATACCCKG-3′ and 1193R: 5′-ACGTCATCCCCACCTTCC-3′) (26, 45, 46). PCR was performed on ABI GeneAmp® 9700 (Applied Biosystems, Foster City, USA) with the following conditions: 2.5 mM dNTPs, 5 μM forward and reverse primers, and 0.4 μL FastPfu polymerase. The cycling process proceeded as follows: 3 min for 95 °C, 27 cycles at the same temperature for 30 s, annealing at 55 °C for 30 s, 72 °C for 15 s, and then extension at 72 °C for 10 min.

### Illumina MiSeq sequencing and data analysis

PCR amplification products of the V5–V7 region of fungal 18S and bacterial 16S rRNA genes were sequenced by Illumina MiSeq sequencer. The Illumina raw sequences were filtered by QIIME (47). Specific barcodes were set to assign the different agarwood samples (48). Chimeric sequences were checked and removed. The taxonomic levels of the bacteria and fungi in this study depended on the OTUs (Operational taxonomic units) at 97% similarity, which was defined on the basis of trimmed sequences. These OTUs were assigned based on the Silva database (http://www.arb-silva.de). The abundance of each OTU in tables described the level of fungal and bacterial phylotypes in different samples. To determine the most representative microbial community, mitochondrial and chloroplast OTUs, as well as those with < 0.1% abundance in different samples, were filtered manually (49). Meanwhile, OTUs were rarefied to the lowest sequence number. The simplified OTU tables were used for the following analysis. Rarefaction curves, Chao and Shannon indices, Venn diagrams, community barplots at different taxonomic levels, and community heatmap analysis were employed to analyze the microbial community structure of the agarwood samples. Bacterial functional profile was used to compare the gene function at different levels derived from the variational microbial community abundance. The raw sequences of all samples reported in this study have been submitted to the National Center for Biotechnology Information Short Read Archive (accession no. SRP127004) under BioProject PRJNA422987.

### Statistical analysis

The results were reported as means ± SD. Data were analyzed using IBM SPSS Statistics 20. Statistically significant data were further analyzed using one-way analysis. Significant difference was considered at *p* < 0.05.

## ACKNOWLEDGMENTS

The authors would like to thank Mr. Zong-Miao Ding (Chairman of Hainan Xiangshu Aloes Industry Co., LTD.) who provided the agarwood samples generously.

## AUTHOR CONTRUBUTIONS

Q.G. Mo completed the experiments via culture-independent methods, interpreted data and developed a manuscript draft. C.Y. Fan and H.Y. Fu completed the experiments via culture-dependent methods. G. Zhou supervised the collection of agarwood samples. Y.W. Wang supervised the whole research and rewrote this manuscript.

## ETHICAL APPROVAL

This article does not contain any studies with human participants or animals performed by any of the authors.

